# The Y chromosome gene KDM5D restrains CD8+ T cell antitumor immunity through TCR and cholesterol-exhaustion programs

**DOI:** 10.64898/2026.07.23.740424

**Authors:** Jiexi Li, Chi Yin Ching, Aviad Ben-Shmuel, Paulino Tallón de Lara, Jingjing Liu, Jiang Shan, Chaohao Li, Ziyuan Zhang, William Huaiyang Wu, Max Slotnik, Xiaotian Wang, Raul Caballero Montes, Abhinav K. Jain, Nicholas Hornstein, Fadl Zeineddine, Mohammad Zeineddine, Scott Eric Woodman, Natividad Roberto Fuentes, Denise J. Spring, John Paul Shen, Scott Kopetz, Ronald A. DePinho

## Abstract

Sex differences in immunity shape cancer risk, autoimmunity, and responses to immunotherapy, yet the sex-chromosome genes that regulate antitumor T cell function remain incompletely defined. Here, we identify the Y chromosome-encoded KDM5D histone demethylase as a male-specific suppressor of CD8+ T cell antitumor immunity. In murine colorectal cancer (CRC) models, male CD8+ T cells displayed reduced cytokine production, proliferation, cytotoxicity, TCRβ abundance, and proximal TCR signaling relative to female CD8+ T cells. CRISPR-RNP-mediated KDM5D depletion in male CD8+ T cells enhanced effector function, increased TCRβ expression, augmented TCR signaling, and improved tumor control after adoptive transfer. Transcriptomic and functional analyses further linked KDM5D to cholesterol biosynthesis and exhaustion-associated programs, with KDM5D depletion reducing SREBP2/XBP1-associated cholesterol and exhaustion signatures. Correspondingly, human CRC single-cell analyses supported the clinical relevance of this axis, showing enrichment of exhausted and cholesterol-associated CD8+ T cell states in male tumors. Pharmacologic inhibition of cholesterol biosynthesis with lovastatin partially attenuated select exhaustion-associated markers in male CD8+ T cells and delayed tumor growth in vivo. Together, these findings define KDM5D as a sex chromosome-encoded regulator of male CD8+ T cell dysfunction and point to cholesterol-exhaustion programs as a potential therapeutic vulnerability in male CRC.

## Introduction

Sex differences in cancers are well established, with males showing higher incidence and mortality across the majority of non-reproductive cancers, whereas females show increased responsiveness to cancer immunotherapy and susceptibility to autoimmune diseases (Klein and Flanagan 2016). This sexual dimorphism is especially evident in colorectal cancer (CRC) across tumor biology, immunity, and clinical outcomes (Baraibar et al. 2023). In CRC, sex differences extend beyond tumor epidemiology to tumor-immune profiles, where multiple studies report different levels of intratumoral lymphocyte infiltration (CD8+, CD4+, and Th2 cells) as well as distinct cytokine and myeloid programs, which track with survival (Ray et al. 2020). CD8+ cytotoxic T cells are central mediators of antitumor immunity, and emerging evidence indicates that their differentiation, metabolic state, and exhaustion programs differ by sex (Kwon et al. 2022; Yang et al. 2022). However, the genetic mechanisms underlying these T cell differences remain incompletely defined.

While sex hormones clearly influence immune function (Klein and Flanagan 2016), sex chromosome genes may represent an underexplored source of sex-specific immune regulation that operates coordinately with hormones. Our previous work demonstrated that the Y chromosome gene KDM5D promotes aggressive male CRC through cancer cell-intrinsic mechanisms, including enhancement of metastatic competence and repression of antigen processing and presentation (Li et al. 2023). In this context, we showed that KDM5D encodes a histone demethylase that removes methyl groups from H3K4 to repress target gene expression, such as cell adhesion genes, including *Amot*, and associates with the Sin3-HDAC complex via SAP18 to alter the super-enhancer that regulates MHC-I gene expression, leading to decreased antigen processing and presentation in cancer cells (Li et al. 2023). Pointing to a potential regulatory role in T cells, a genome-scale CRISPR screen in male CD8+ T cells identified KDM5D as a top-ranked suppressor of T cell function (Kumar et al. 2021). Together, these findings raised the possibility that KDM5D may operate in parallel in malignant epithelial cells and male CD8+ T cells, thereby reinforcing male-biased immune escape through both cancer cell-intrinsic and immune cell-intrinsic mechanisms.

CD8+ T cell function is governed by the integration of TCR signal strength, co-stimulation, metabolic fitness, and chronic antigen exposure. TCR engagement initiates proximal phosphorylation events involving LCK, ZAP70, LAT, PLCγ1, and ERK, ultimately driving calcium flux, transcription factor activation, cytokine production, and cytotoxicity (Gaud et al. 2018). Sex-biased regulation of TCR signaling has been documented in autoimmune settings, including androgen receptor (AR) signaling induction of negative TCR signaling regulators such as PTPN22, which raises the activation threshold in male CD8+ T cells and partially accounts for lower susceptibility of men to autoimmune diseases (Stanford and Bottini 2014; Lee et al. 2024). Whether analogous sex differences in TCR signaling in the TME, and whether sex chromosome genes other than those encoding hormones are responsible, remain important open questions.

T cell metabolism provides a second layer of T cell regulation (Madden and Rathmell 2021). Female CD8+ T cells exhibit greater mitochondrial respiratory capacity and fitness, whereas male CD8+ T cells are more susceptible to mitochondrial stress, reactive oxygen species, lipid peroxidation, and endoplasmic reticulum (ER) stress (Lee et al. 2025; Madi et al. 2026). Cholesterol homeostasis is particularly relevant in that excessive intracellular cholesterol can promote ER stress, XBP1 activation, and CD8+ T cell exhaustion (Ma et al. 2019; Shuwen et al. 2023; Hu et al. 2024). Thus, the balance of cholesterol synthesis and stress signaling may be a critical determinant of sex-biased CD8+ T cell states in the TME.

Here, we identify KDM5D as a male-specific, cell-intrinsic suppressor of CD8+ T cell effector function in CRC models. Experimental and translational computational data support a framework in which KDM5D attenuates TCR proximal signaling while promoting cholesterol-associated exhaustion programs. These findings provide a genetic and metabolic framework for sex-informed cancer immunotherapy enhancement strategies.

## Results

### KDM5D restrains male CD8+ T cell effector function

We previously reported that KRAS-mutant CRC exhibits a sex-biased progression involving both cancer cell-intrinsic and immune evasion mechanisms (Li et al. 2023). To extend this work to a faithful model of the most common form of CRC, we utilized an established and well-characterized genetically engineered mouse model that incorporates APC and p53 loss, KRAS-G12D expression (Boutin et al. 2017), and chromosomal instability (CIN) induced through telomere dysfunction and telomerase reactivation (LaBella et al. 2024). This model, hereafter termed CIN+ iT-KAP, mirrors well the features of the most common CRC subtype, microsatellite-stable (MSS)-CRC, which features an immune-cold tumor microenvironment and poor responsiveness to immunotherapy (Xia et al. 2016). Orthotopic transplantation of male or female CIN+ iT-KAP organoids into sex-matched immunocompetent C57BL/6 or immunodeficient NSG mice revealed pronounced sex-dependent differences in tumor growth exclusively in the immunocompetent setting (**Fig. 1A**), indicating that adaptive immunity is involved in the observed sex bias in tumor progression. Although tumor organoids were implanted into sex-matched hosts and therefore do not formally distinguish host-from tumor-intrinsic contributions, our findings are consistent with prior studies implanting identical tumor cell lines into both male and female hosts. These studies similarly demonstrated enhanced CD8+ T cell-dependent tumor control in females despite identical tumor cells, supporting intrinsic sex differences in CD8+ T cell function (Dakup et al. 2020; Ray et al. 2020; Kwon et al. 2022; Song et al. 2022). Together, these data reinforce the role of host immunity in shaping sex differences in tumor progression. To test whether CD8+ T cells harbor cell-intrinsic, sex-biased functional properties beyond those attributable to circulating sex hormones, we isolated CD8+ T cells from male and female OT-1 littermates and activated them in vitro with anti-CD3/CD28 beads and recombinant IL-2. Female CD8+ T cells produced higher IFN-γ and Granzyme B and showed increased Ki67-positivity relative to male CD8+ T cells (**Fig. 1B,C)**. In an antigen-specific cell killing assay, we used the CMT93-KRAS-G12D-OVA CRC model as target cells, as this cell line is sex-neutral due to spontaneous loss of the Y chromosome (Hussein Azawi et al. 2018), expresses KRAS-G12D to recapitulate the impaired immunogenicity of human CRC, and stably expresses ovalbumin as a model antigen. Co-culture of these target cells with male or female OT-1 CD8+ T cells showed superior cytotoxicity by female T cells at different effector-to-target ratios (**Fig. 1D**). These results indicated that male and female CD8+ T cells differ intrinsically in effector fitness under controlled culture conditions with minimized hormonal influences.

**Fig. 1.**
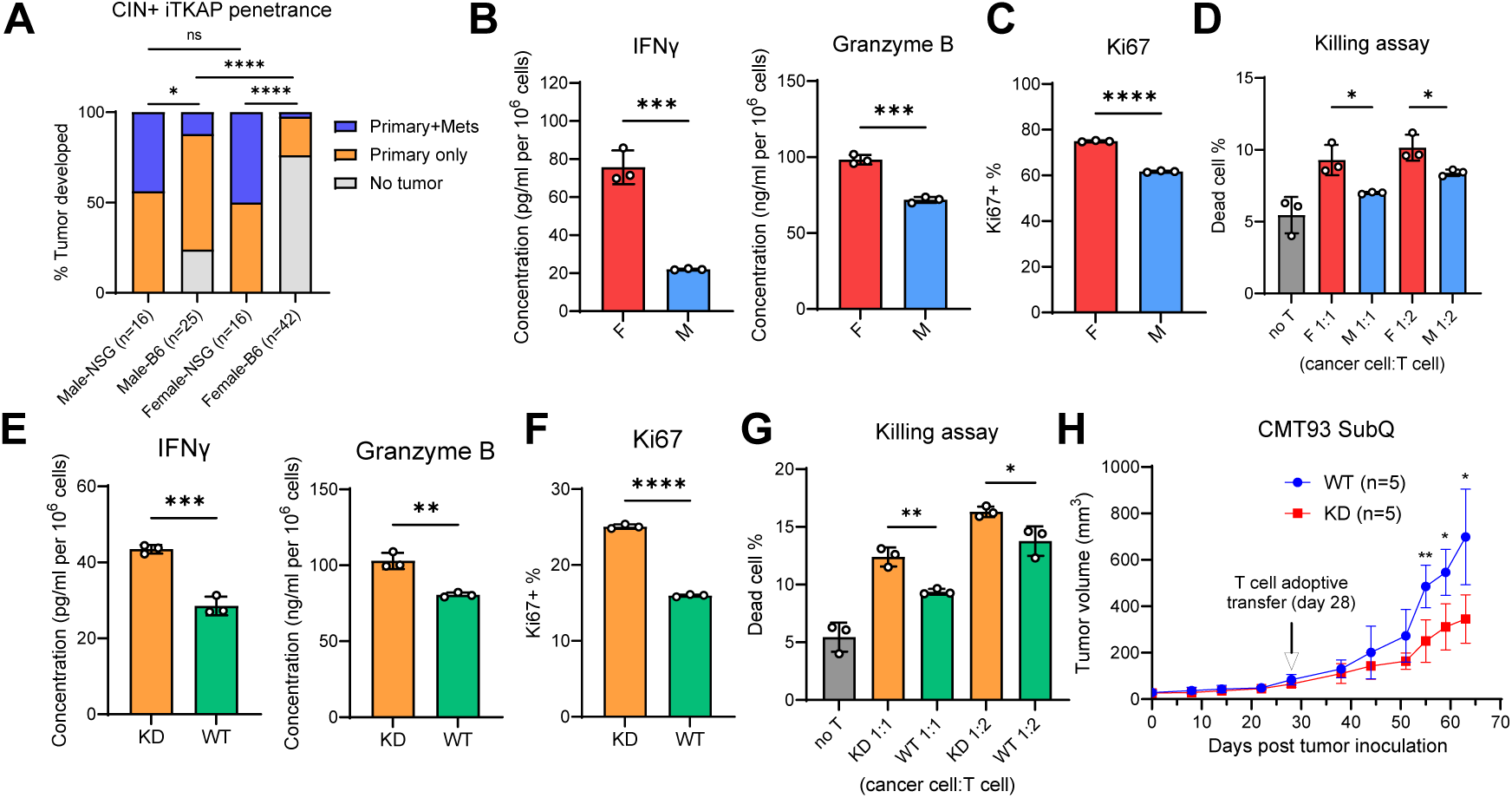
KDM5D restrains male CD8+ T cell effector function. **A**. Growth capability of CIN+ iT-KAP organoid in B6 versus NSG mice of both sexes (Fisher’s Exact test). Mets: metastasis. **B**. ELISA of cytokines from the same number of male and female OT-1 CD8+ T cells. **C**. Flow cytometry of Ki67. **D**. OT-1 T cell killing assay. Cancer cells: CMT93-KRAS-G12D-OVA. **E**. ELISA of cytokines from the same number of male OT-1 CD8+ T cells with or without KDM5D KD. **F**. Flow cytometry of Ki67. **G**. OT-1 T cell killing assay. Cancer cells: CMT93-KRAS-G12D-OVA. **H**. Subcutaneous (SubQ) tumor growth of CMT93-KRAS-G12D-OVA in nude mice with adoptive transfer of male OT-1 CD8+ T cells with or without KDM5D KD. For **B-H**, data are mean value ± s.d.; *P* was derived with two-tailed unpaired t-test.

Because our previous studies implicated KDM5D in colorectal cancer cell biology (Li et al. 2023) and an independent CRISPR screen identified KDM5D as a top candidate negative regulator of male CD8+ T cell function (Kumar et al. 2021), we next depleted KDM5D in male OT-1 CD8+ T cells using CRISPR-ribonucleoprotein (RNP) electroporation. Because editing generated a mixed population rather than clonal knockout (KO) cells, we refer to this condition as KDM5D knockdown/depletion (KD). Control cells underwent electroporation but received only sgRNA without SpCas9. Significant KDM5D depletion (> 50%) was confirmed by qPCR in cultured cells (**Supplemental Fig. 1A;** representative data; similar KD efficiency was observed in all other experiments). KDM5D depletion increased IFN-γ and Granzyme B secretion, enhanced proliferation (Ki67 positivity), and improved killing of CMT93-KRAS-G12D-OVA target cells (**Fig. 1E-G**). In adoptive-transfer experiments using CMT93-KRAS-G12D-OVA tumors in male nude mice, KDM5D-depleted OT-1 T cells delayed tumor growth relative to control male OT-1 cells (**Fig. 1H; Supplemental Fig. 1B**).

We further assessed transferred OT-1 cells within tumors in an immune-competent context using the CD45.1/CD45.2 tracking system, in which CD45.1 host mice with CIN+ iT-KAP OVA tumors were adoptively transferred with male OT-1 CD8+ T cells (CD45.2), either KD or WT for KDM5D (**Supplemental Fig. 1C**). Although transferred T cells produced only transient tumor control, analysis at a time point when tumor sizes were comparable (16 days post T cell transfer) revealed that infiltrating OT-1 CD8+ T cells (CD3+ CD8+ CD45.2+; **Supplemental Fig. 1D,E**) with KDM5D depletion contain a higher proportion of Perforin+ Granzyme B+ cytotoxic CD8+ T cells and lower expression of exhaustion markers including TIM-3 and TOX with trends toward reduced LAG-3 and PD-1 (**Supplemental Fig. 1F-J**).

Correspondingly, the correlation of KDM5D and exhaustion is evident in mouse CRC single-cell datasets of spontaneous male iKAP tumors, which show highest KDM5D expression in the exhausted CD8+ T cells (**Supplemental Fig. 2A**); similarly, male CRC patient tumors exhibit the lowest KDM5D expression in cytotoxic CD8+ T cells and highest in exhausted CD8+ T cells (**Supplemental Fig. 2B,C**). Specifically, within the CD8+ T cell compartment, male cells exhibited more exhausted T cells, higher expression of immune checkpoint/exhaustion genes, including *PDCD1* (PD1), *CTLA4*, *LAG3*, *HAVCR2* (TIM3), *TIGIT* and *TOX*, and lower levels of cytotoxic markers, such as *PRF1* (Perforin), *IFNG* (IFN-γ) and *TNF* (TNF-alpha) (**Supplemental Fig. 2D,E**). Together, these data support a model in which KDM5D constrains male CD8+ T cell effector function and promotes exhaustion-associated states in the tumor setting.

### KDM5D restrains TCR abundance and proximal signaling

To define the mechanisms through which KDM5D suppresses male CD8+ T cell function, we compared the transcriptomes of male versus female CD8+ T cells and of KDM5D-depleted versus control male CD8+ T cells. Pathway analyses of genes increased in both female cells and KDM5D-depleted male cells identified consistent enrichment of CDC42, RAC3, and RHOJ GTPase cycle programs (**Fig. 2A**). Given that CDC42 Rho-family GTPases are central regulators of T cell receptor (TCR) signaling and T cell activation, and coordinate cytoskeletal remodeling, TCR microcluster organization, and immunological synapse formation (Tskvitaria-Fuller et al. 2006; Tybulewicz and Henderson 2009; Gaud et al. 2018), we examined whether KDM5D influences TCR abundance signaling.

**Fig. 2.**
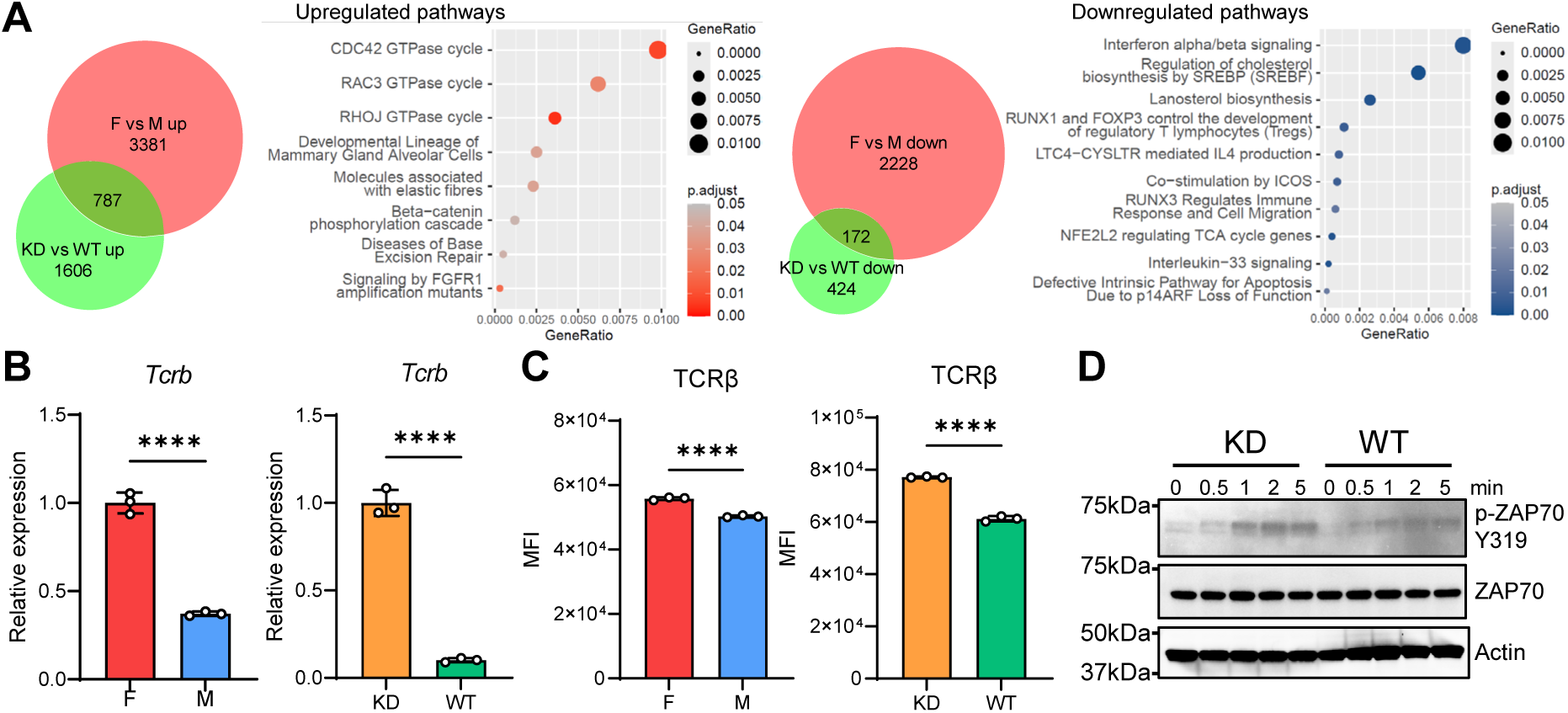
KDM5D restrains TCR abundance and proximal signaling. **A**. Pathway analysis of the differentially expressed genes. **B**. qPCR of *Tcrb* in OT-1 T cells. **C**. Flow cytometry of cell surface TCRβ. **D**. Western blot of phos-ZAP70 Y319, total ZAP70 and Actin in restimulated cells. For **B** and **C**, data are mean value ± s.d.; *P* was derived with two-tailed unpaired t-test.

Indeed, TCRβ mRNA and protein levels were lower in male than in female OT-1 CD8+ T cells and were upregulated after KDM5D depletion in male OT-1 cells (**Fig. 2B,C; Supplemental Fig. 3A**). Similar sex differences were observed in polyclonal C57BL/6 CD8+ T cells (**Supplemental Fig. 3B**), supporting the relevance of this finding beyond the OT-1 system. It is worth noting that, in the OT-1 system, while the TCRβ transgene is integrated into chromosome 3, it remains under the control of the endogenous Eβ, located approximately 5-7.5 kb downstream of the T Cell Receptor Beta Constant 2 (*Trbc2*) gene on chromosome 6 (Rodriguez-Caparros et al. 2020). CUT&RUN profiling demonstrated increased H3K4me1 and H3K4me2 signal at the *Trbc2* regulatory region after KDM5D depletion, consistent with enhancer-associated chromatin activity; moreover, the absence of H3K4me3 (a promoter-specific mark) further supports the enhancer identity of this region (**Supplemental Fig. 3C**). To investigate whether KDM5D’s enzymatic domain is required for regulating TCRβ expression, we overexpressed WT or enzymatic-dead (ED) mouse KDM5D in female OT-1 CD8+ T cells. Only WT, but not ED, KDM5D robustly repressed TCRβ expression (**Supplemental Fig. 3D**). Together, these data suggest that KDM5D depletion elevated Eβ activity and TCRβ expression through an enzymatic function-dependent mechanism.

Consistent with these changes in TCRβ level, female CD8+ T cells and KDM5D-depleted male CD8+ T cells showed elevated phosphorylation of key TCR signaling molecules, including LCK, ZAP70, LAT, PLCγ1 and ERK (**Supplemental Fig. 3E**). Following antigen-specific restimulation with H2Kb/SIINFEKL tetramer, KDM5D-depleted male OT-1 CD8+ T cells showed more rapid and stronger ZAP70 phosphorylation than control cells, suggesting faster activation kinetics in response to antigen engagement (**Fig. 2D**). Together, these findings support the view that KDM5D serves as an epigenetic regulator of TCR abundance and proximal signaling activation in male CD8+ T cells.

### KDM5D regulates cholesterol homeostasis and exhaustion-associated programs in male CD8+ T cells

Transcriptomic analysis also revealed that cholesterol biosynthesis programs, including SREBP/SREBF-regulated pathways and lanosterol biosynthesis, were significantly enriched in male relative to female CD8+ T cells and reduced by KDM5D depletion (**Fig. 2A**). Correspondingly, cholesterol biosynthesis genes, including the master transcription factor *Srebf2* (encoding SREBP2) (Brown and Goldstein 1997; Madison 2016), *Hmgcs1*, *Hmgcr*, *Fdps*, *Fdft1*, *Sqle*, and *Lss* were higher in male CD8+ T cells and reduced by KDM5D depletion (**Fig. 3A**). A similar pattern was observed in CRC patients, where the cholesterol biosynthesis pathway and the expression of cholesterol synthesis genes were more active in male CD8+ T cells than in female cells (**Fig. 3B; Supplemental Fig. 4A**).

**Fig. 3.**
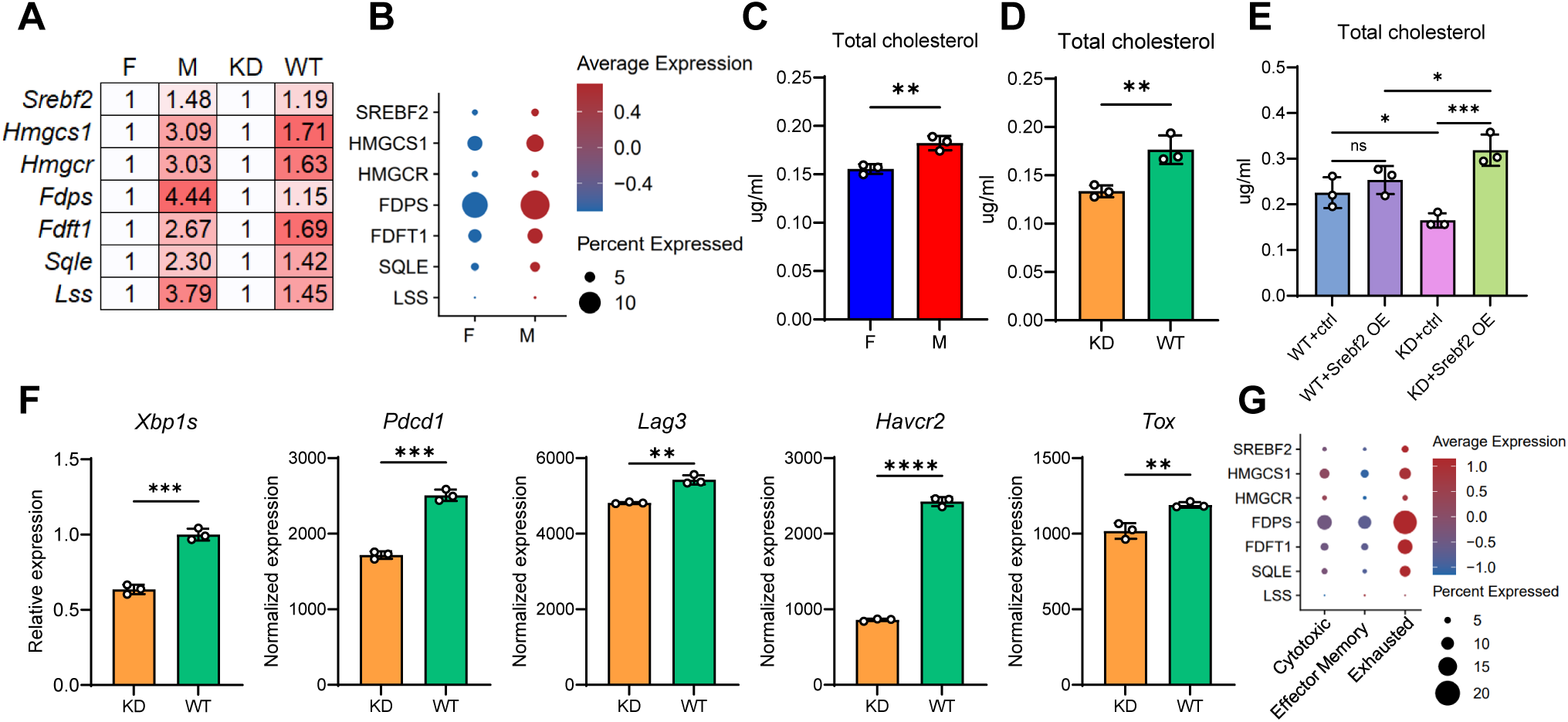
KDM5D regulates cholesterol homeostasis and exhaustion-associated programs in male CD8+ T cells. **A**. Fold change of genes in female versus male CD8+ T cells and male T cells with KDM5D KD versus WT from RNAseq data. **B**. Expression of indicated genes in patient scRNAseq. **C-E**. Total cholesterol levels of the corresponding cells. **F**. qPCR of *Xpb1s* and RNAseq of immune checkpoint genes. **G**. Expression of indicated genes in patient scRNAseq. For **C-F**, data are mean value ± s.d.; *P* was derived with two-tailed unpaired t-test (**C**, **D** and **F**) and one-way ANOVA with multiple comparisons (**E**).

Male CD8+ T cells contained higher total cholesterol levels than female cells, and KDM5D depletion reduced total cholesterol in male CD8+ T cells (**Fig. 3C,D**). Overexpression of *Srebf2* in KDM5D-depleted T cells restored total cholesterol levels, placing SREBP2 downstream of, or parallel to, KDM5D-dependent cholesterol regulation (**Fig. 3E; Supplemental Fig. 4B,C**). Excessive cholesterol is known to promote CD8+ T cell exhaustion by triggering ER stress and activating the unfolded protein response, particularly through *Xbp1* splicing, which in turn increases immune checkpoint gene expression, including PD-1, 2B4, TIM-3, and LAG-3 (Ma et al. 2019). We additionally observed that KDM5D depletion attenuates this program in male CD8+ T cells, reducing *Xbp1* splicing (*Xbp1s*), *Tox*, and checkpoint receptor expression (**Fig. 3F**). Using an in vitro chronic-stimulation model with anti-CD3/CD28 beads every 3 days versus single stimulation, representing acute activation (Lacalle et al. 2025) (**Supplemental Fig. 4D**), we confirmed successful induction of exhaustion markers in chronically stimulated cells with upregulation of *Pdcd1* (PD-1), *Havcr2* (TIM-3), and *Lag3* genes as well as *Srebf2* and *Xbp1s* (**Supplemental Fig. 4E**). In chronically stimulated cells, KDM5D depletion attenuated induction of *Pdcd1*, *Havcr2*, *Lag3*, *Srebf2* (with trend though not significant), and *Xbp1s* relative to WT controls (**Supplemental Fig. 4E**), strongly indicating that KDM5D depletion can impede the transition of male CD8+ T cells toward a more exhausted state. Furthermore, human and mouse single-cell transcriptomic analyses support these experimental observations. Cholesterol-associated programs were enriched in the exhausted CD8+ T cells compared with cytotoxic CD8+ T cells; and among the exhausted CD8+ T cells, the cholesterol homeostasis pathway was more active in males than female cells (**Fig. 3G; Supplemental Fig. 4F-H**).

Given the epigenetic regulatory activity of KDM5D, we hypothesized that KDM5D may regulate cholesterol metabolism-associated genes on the epigenetic level. Indeed, only WT, not ED, KDM5D induced the expression of immune checkpoint genes, as well as *Tox*, *Srebf2*, and *Xbp1s* (**Supplemental Fig. 5A**). *SREBF2* gene expression has been reported to be repressed by sirtuin (Sirt) 6. Sirt6 is recruited by forkhead box O (FoxO)3 to the *SREBF2* gene promoter, where it deacetylates histone H3 at lysines 9 and 56, thereby establishing a repressive chromatin state (Tao et al. 2013). This raises the possibility that KDM5D might upregulate SREBP2 expression by suppressing FOXO3 expression. In CD8+ T cells, CUT&RUN profiling showed that KDM5D KD increased the levels of H3K4me2/3 and H3K27ac at the *Foxo3* gene promoter and enhancer regions (**Supplemental Fig. 5B**). Consistent with the histone modification changes, *Foxo3* expression was also upregulated in KDM5D-KD T cells (**Supplemental Fig. 5C**). Moreover, *Foxo3* expression is higher in female CD8+ T cells than in male cells (**Supplemental Fig. 5D**). Similarly, in CD8+ T cells from CRC patient tumors, *FOXO3* expression was higher in females than in males and was also lowest in exhausted CD8+ T cells (**Supplemental Fig. 5E**). Together, these data support the view that the KDM5D–FOXO3–SREBP2 axis may contribute to sex differences in CD8+ T cell function.

### Lovastatin attenuates exhaustion-associated markers and delays male CRC tumor growth

Given that KDM5D is associated with increased cholesterol biosynthesis and exhaustion-associated programs in male CD8+ T cells, we asked whether pharmacologic inhibition of cholesterol biosynthesis could improve male CD8+ T cell function (Kansal et al. 2023). In vitro lovastatin treatment (100nM) reduced *Pdcd1* and *Lag3* expression in male CD8+ T cells at 48 hours; and, notably, no such effect was observed in female cells (**Fig. 4A,B**). Moreover, lovastatin did not affect L2-3 male iKAP metastatic cancer cell proliferation in vitro (**Fig. 4C**), suggesting it exerts minimal direct cancer cell-intrinsic effects.

**Fig. 4.**
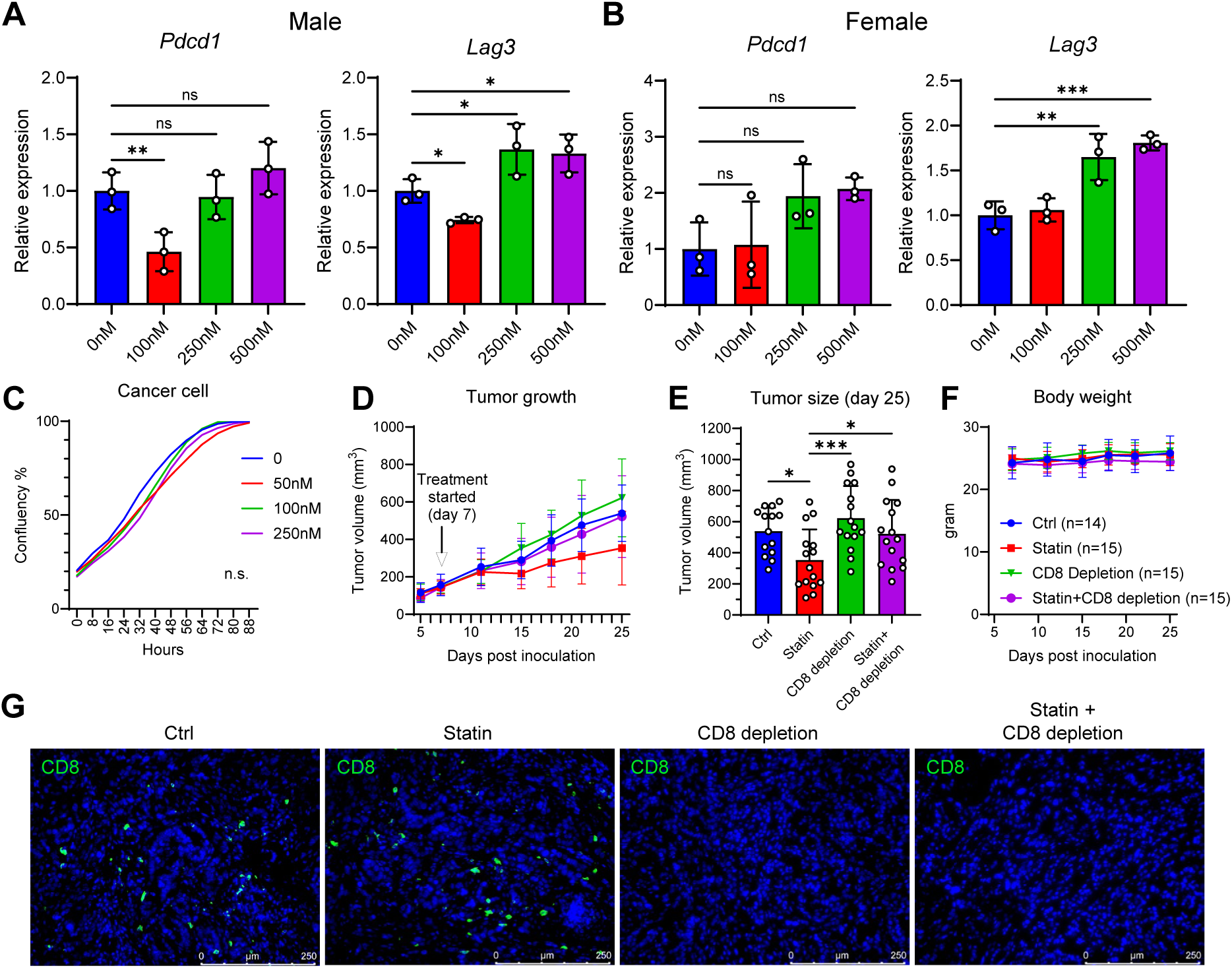
Lovastatin attenuates exhaustion-associated markers and delays male CRC tumor growth. **A**, **B**. qPCR of *Pdcd1* and *Lag3* in male and female CD8+ T cells treated with different dosages of lovastatin for 48 hours. **C**. IncuCyte proliferation assay of male iKAP cancer cell L2-3 after one-time treatment of lovastatin at the indicated dosages (two-tailed Mann-Whitney test). **D**. Growth curve of male CIN+ iT-KAP subcutaneous tumors. **E**. Tumor size of the mice in **D** on day 25 after tumor inoculation (only significant *P* values were marked). **F**. Weight of the mice in **D**. **G**. Immunofluorescence staining of CD8 (green) and nucleus (DAPI) using tumors from **D**. For **A**, **B** and **E**, data are mean value ± s.d.; *P* was derived with one-way ANOVA with multiple comparisons.

In male mice bearing CIN+ iT-KAP tumors, lovastatin treatment delayed tumor growth without measurable weight loss, and CD8+ cell depletion abolished the treatment effect (**Fig. 4D-G**). Together with prior reports linking statin use to sex-biased CRC risk reduction and potential synergy with immune checkpoint blockade (Clancy et al. 2013; Shah et al. 2024), these data support cholesterol biosynthesis as a tractable pathway for modulating male CD8+ T cell dysfunction.

## Discussion

In this study, we identify the Y chromosome gene KDM5D as a male-specific regulator of CD8+ T cell function, highlighting the role of sex chromosome genes in immune sexual dimorphism in cancer. Mechanistically, we show that male CD8+ T cells display reduced effector function, diminished TCR expression, attenuated proximal TCR signaling, elevated cholesterol biosynthesis programs, and enhanced exhaustion-associated states. KDM5D depletion partially reverses these phenotypes, supporting a model in which KDM5D integrates epigenetic and metabolic mechanisms to restrain male CD8+ T cell antitumor activity.

A central finding is the role of KDM5D in dampening the TCR signaling axis, defining a novel dimension in sex-specific CD8+ T cell regulation. KDM5D/sex-dependent transcriptomic profiles pointed to the CDC42 GTPase pathway, a downstream branch of TCR signaling, prompting integrated omic and functional analyses. The convergence of transcriptomic pathway analyses, TCRβ expression, and proximal phosphorylation assays supports a functional relationship between KDM5D and TCR activation capacity. These findings align with the capacity of CDC42 to coordinate T cell microvilli formation, TCR microcluster organization, and immunological synapse assembly, thereby facilitating antigen recognition, TCR activation, and antigen-presenting cell engagement (Soh et al. 2025). On the epigenetic level, KDM5D is a known histone demethylase for H3K4me2/3 marks and regulates histone deacetylation via interaction with the Sin3-HDAC complex (Li et al. 2023). Here, we further show that KDM5D influences H3K4me1, implicating engagement of additional histone-modifying complexes in the context of T cells. Although KDM5D-dependent regulation of TCRβ expression was observed in both OT-1 and polyclonal C57BL/6 CD8+ T cells, the chromatin analyses were performed following acute KDM5D depletion in vitro. Thus, future studies using physiological genetic models will be valuable to further define how KDM5D establishes TCR regulatory programs during endogenous T-cell development and activation.

A second key finding is the link between KDM5D and cholesterol-associated exhaustion. Excessive cholesterol has been shown to induce ER stress and XBP1-dependent exhaustion programs in CD8+ T cells (Ma et al. 2019). Our data indicate that male CD8+ T cells have higher cholesterol and cholesterol biosynthesis gene expression, both of which are reduced by KDM5D depletion. The restoration of cholesterol levels by SREBP2 overexpression supports the relevance of SREBP2-dependent biosynthesis, while chromatin and expression changes of FOXO3 suggest a potential upstream epigenetic node. These data establish a framework in which KDM5D promotes a cholesterol-rich, exhaustion-prone state in male CD8+ T cells.

From a translational standpoint, patient single-cell analyses corroborated the highest KDM5D expression and cholesterol pathway activity in exhausted CD8+ T cells in male CRC patient tumors, providing proof-of-principle clinical relevance. In addition, our findings point to a potential therapeutic strategy for the management of male CRC via repurposing statins. We show that statins could suppress immune checkpoint gene expression in male CD8+ T cells in vitro and effectively delay CRC tumor growth in male mouse models in a CD8+ T cell-dependent manner. Previous studies have reported the effects of sex hormones in CD8+ T cells. AR signaling in CD8+ T cells drives a sex-biased exhaustion program impairing antitumor immunity in males, an effect reversed by AR ablation and further enhanced by combination with immune checkpoint blockade (Kwon et al. 2022; Yang et al. 2022). Given that cholesterol is a key precursor for androgen biosynthesis, it is tempting to speculate that statin-mediated HMGCR inhibition could also reduce androgen production to enhance CD8+ T cell effector function in the tumor microenvironment. Beyond this proposed mechanism, statins have pleiotropic antitumor effects on other immune cells, including enhancing dendritic cell antigen presentation and repolarizing M2-like toward M1-like macrophages (Khan et al. 2009; Mira et al. 2013; Al Dujaily et al. 2020). In addition, cholesterol-independent effects of statins, including HDAC inhibition (Singh et al. 2016), may enhance chromatin accessibility at TCRβ gene regulatory loci. Nevertheless, these effects vary with statin type, dose, and tumor context, likely accounting for the inconsistency observed across clinical studies. Therefore, future work should test whether the antitumor benefit of statins depends on KDM5D or SREBP2/XBP1 signaling, and whether it synergizes with immune checkpoint blockade in male CRC.

While our data are confined to the tumor context, it is tempting to speculate that sex differences in CD8+ T cell biology may extend to other disease settings. These include female-biased autoimmune diseases such as multiple sclerosis, systemic lupus erythematosus, and rheumatoid arthritis, which have been linked to aberrant or heightened CD8+ T cell activity (Carvalheiro et al. 2013; Deng et al. 2019). Direct testing in relevant disease models will be needed to evaluate these possibilities.

In conclusion, our study demonstrates that the Y chromosome gene KDM5D is a key regulator of sex differences in CD8+ T cell function in CRC via dual regulation of TCR epigenetic silencing and cholesterol-induced T cell exhaustion (**Supplemental Fig. 6**). These findings elucidate a novel genetic and metabolic mechanism underlying tumor immune sex disparities and support statin repurposing as a potential therapy for male CRC patients. Future studies will focus on validating the clinical efficacy of statin-based sex-specific therapy and developing KDM5D-specific inhibitors, ultimately promoting the development of precision immune therapy for CRC and other malignant tumors.

## Materials and Methods

### Mouse models

CIN+ iT-KAP mice (tet-O-LSL-Kras^G12D^;Apc^fl/fl^;Trp53^fl/fl^;LSL-mTERT;LSL-rtTA-GFP;Villin-creER^T2^) were generated using previously published alleles (Boutin et al. 2017; LaBella et al. 2024). The C57BL/6 (B6), C57BL/6-Tg (TcraTcrb)1100Mjb/J (OT-1) mice, NU/J (nude) mice, and NSG mice were purchased from the Jackson Laboratory.

### Animal experiments

All mouse manipulations were reviewed and approved by The University of Texas MD Anderson Cancer Center’s Institutional Animal Care and Use Committee (IACUC). All animals were maintained in pathogen-free conditions and cared for in accordance with policies and certification of the Association for Assessment and Accreditation of Laboratory Animal Care International (AAALAC International). Housing rooms were maintained on a 12/12 h light/dark cycle (lights on at 06:00, lights off at 18:00) with humidity ranging from 30 to 70%. Temperature ranged from 20 to 24 °C (68 to 76 °F).

For the tumor penetrance assay, normal colonoids from healthy male and female CIN+ iT-KAP mice were treated with 10 mM 4-OHT to induce recombination and mutations, following the method described previously (LaBella et al. 2024). The colonoids were cultured in growth factor-reduced, phenol-free Matrigel (Corning) and media containing Advanced DMEM/F12 (Gibco), 100ng/mL Noggin (SinoBiological, catalog no. 50688-M02H-1), B27 (MCE, catalog no. HY-K3013, 50 ×), N2 (MCE, catalog no. HY-K3012, 100 ×), 1 mM *N*-acetylcysteine (Sigma, catalog no. A9165), 10 mM nicotinamide (Sigma, catalog no. N0636), 10mM HEPES (Gibco), 2mM (1 ×) GlutaMax (Gibco), 100 ug/ml Primocin (InvivoGen), 1 × penicillin/streptomycin (Gibco) with 1 μg/mL doxycycline to induce oncogenic *Kras* G12D expression. For orthotopic implantation, colonoids were digested with TrypLE (Gibco) to remove Matrigel and generate single cell suspension. For each mouse, 0.5 × 10^6^ cells were resuspended in culture media with 20% Matrigel. Mice were anesthetized with isoflurane, and colons were flushed 3 times with sterile PBS. Then, cells were injected into the submucosal layer of either C57BL/6 or NSG male and female mice using a colonoscopy-guided injection (Karl Storz). For *Kras* G12D expression, mice were provided with doxycycline in drinking water (2 mg/ml) throughout the duration of the experiment.

For subcutaneous tumor cell inoculation, 8-week-old mice were anesthetized with isoflurane. Cells were resuspended in Matrigel (Corning) and injected subcutaneously into the right flank using a 27 G needle. Tumor size was estimated with the formula: volume = length × width × width / 2. Tumor size was measured using a digital caliper. For the in vivo killing assay, adoptive transfer of edited CD8+ T cells was done via tail vein injection of T cells resuspended in PBS.

The number of mice per group was decided based on previous experience (Boutin et al. 2017), common practice in the field, animal welfare guidelines, and availability of animals, while minimizing the use of animals in accordance with animal care guidelines from The University of Texas MD Anderson Cancer Center’s IACUC and the National Institutes of Health (NIH).

### Murine T cell isolation and culture

Eight-week-old male OT-1 mice or C57BL/6 mice were euthanized following IACUC protocol and the spleen was removed. The spleen was disrupted in PBS containing 2% FBS. The aggregates and debris were removed by passing the cell suspension through a 70 μm mesh nylon strainer. CD8+ T cells were isolated using EasySep™ Mouse CD8+ T Cell Isolation Kit (STEMCELL) following the manufacturer’s protocol. The isolated cells were stimulated with Dynabeads™ Mouse T-Activator CD3/CD28 for T cell Expansion and Activation (ThermoFisher) at a cell-to-bead ratio of 2:1, and were cultured in complete T cell culture media (RPMI-1640 + 2.05 mM l-Glutamine (Cytiva) supplemented with 10% FBS, 1 × penicillin/streptomycin, 1× sodium pyruvate, 1 × nonessential amino acids, 1× β-mercaptoethanol, and 1× GlutaMAX (all from Gibco)) with 30 U/ml mouse recombinant IL-2 (STEMCELL) (Herda et al. 2021).

### CD8+ T cell gene knockout and overexpression

Editing of OT-I CD8+ T cells (3 days after initial stimulation) was performed by electroporation with RNP composed of Cas9 protein and sgRNA of mouse *Kdm5d* (Mouse Gene Knockout Kit v2, EditCo) using the P3 primary cell nucleofector kit (Lonza). The cells were then cultured for an additional three days in complete T cell culture media with 30 U/ml mouse recombinant IL-2, and the editing efficiency of the gene was confirmed by qPCR analysis.

Overexpression of mouse *Srebf2* was achieved with spin infection of the mixture of lentivirus-containing plasmids (pLV-mCherry/Puro-EF1A-mSrebf2 and empty vector control (VectorBuilder)) and T cells in suspension with polybrene for 30 minutes at 800 g at 32 °C. Lentivirus containing the desired plasmids was produced in HEK293T (ATCC, catalog no. CRL-3216, authenticated by ATCC) by Lipofectamine 2000 (Thermo Fisher Scientific) transfection.

Overexpression of mouse wild-type (WT), enzymatic-dead (ED) KDM5D using previously described plasmids (Li et al. 2023) or control vector (pLenti-EF1a-C-tGFP, OriGene, catalog no. PS100072) was achieved with electroporation described above.

### Cancer cell and organoid line generation and culture

The iKAP cell line was established previously (Li et al. 2023). The CMT93 cell line was purchased from ATCC (catalog no. CCL-223, authenticated by ATCC) and cultured with the recommended conditions. To introduce the expression of ovalbumin, mouse *Kras*-G12D and luciferase reporter, the cells were infected with lentivirus containing mouse *Kras*-G12D (cloned into pLenti6.3/V5-DEST vector) overexpression plasmid (Liao et al. 2019), ovalbumin overexpression plasmid (pLVX-puro-cOVA-IRES-BFP, Addgene Plasmid #135074) and RFP-luciferase plasmid (System Biosciences, catalog no. BLIV502MC-1). Infected cells were selected with Blasticidin, followed by flow sorting for BFP-positive, RFP-positive, and H2Kb-SIINFEKL-positive cells simultaneously to confirm *Kras*-G12D, ovalbumin, and luciferase co-expression and successful ovalbumin antigen processing and presentation. H2Kb-SIINFEKL antibody (BioLegend, catalog no. 141606) and IgG isotype control (BioLegend, catalog no. 400120) were used for staining.

For CIN+ iT-KAP tumor organoids, tumors from a male CIN+ iT-KAP mouse were collected and dissociated with Mouse Tumor Dissociation Kit (Miltenyi Biotec). Cancer cells were cultured in Matrigel and the medium described in the “Animal Experiments” section.

To express ovalbumin in organoids, organoids were dissociated with TrypLE (Gibco) for 5 min at 37 °C, washed with Hank’s Balanced Salt Solution (Gibco) with FBS, and spun down at 400 g. The organoid pellets were resuspended in Advanced DMEM/F12 (Gibco) medium supplemented with 10 μM Y-27632 (EMD Millipore) and mixed with lentivirus and polybrene in an ultra-low attachment plate (Corning), then spun at 600 g for 60 min at 32 °C. The cells were incubated at 37 °C for 4 h for recovery and then seeded in Matrigel (Corning). The successfully infected organoids expressing ovalbumin were sorted using BFP and H2Kb-SIINFEKL antibody described above.

### ELISA

An equal number of cells per condition was plated the day before conditioned media collection. Cell number was counted again right before the media collection. The media was spun down, and the supernatant was collected and analyzed using the Mouse IFN gamma ELISA Kit (Abcam, catalog no. ab282874) and Mouse Granzyme B ELISA Kit (Abcam, catalog no. ab238265) following the manufacturers’ protocols.

### Flow cytometry

The recommended protocol, BioLegend Cell Surface Flow Cytometry Staining Protocol, was followed. Briefly, the live cells were washed in Cell Staining Buffer (BioLegend), centrifuged at 350 g for 5 min at 4 °C, stained with TruStain FcX Plus anti-mouse CD16/32 antibody (BioLegend, catalog no. 156604, 1:50) and Ghost Dye UV450 viability stain (TONBO, catalog no. 13-0868-T100, 1:1000) for 30 min at 4 °C in the dark, then stained for 20 min on ice in the dark with TCR beta antibody (BioLegend, catalog no. 109211). Samples were acquired on a Cytek Aurora spectral flow cytometer and analyzed in FlowJo.

To sort out the tumor-infiltrating CD8+ T cells from CMT93-KRAS-G12D-OVA tumors, tumors were dissociated with Mouse Tumor Dissociation Kit (Miltenyi Biotec) stained for CD45-APC (Invitrogen, catalog no. 17-0451-82), CD3-BV510 (BioLegend, catalog no. 100233), CD8-BV785 (BioLegend, catalog no. 100749) and SYTOX Green Dead Cell Stain (Invitrogen).

For Ki67 staining, cells were harvested, pelleted, and fixed by dropwise addition of ice-cold 70% ethanol with continuous vortexing, followed by incubation at -20 °C for a minimum of 2 h. Fixed cells were washed twice in staining buffer and stained with anti-Ki-67 antibody (BioLegend, catalog no. 151209) for 30 min at room temperature. Samples were then washed with staining buffer and proceeded to acquisition on a flow cytometer.

### T cell killing assay

CMT93-KRAS-G12D-OVA cells were treated with 10 ng/ml recombinant mouse IFN-γ protein (Abcam, catalog no. ab9922) for 24 h. IFN-γ was then washed off, the cells were detached and counted, and the activated CD8+ T cells were added at the indicated cancer cell-to-T cell ratios. The next day, all cells were collected and stained with SYTOX Green Dead Cell Stain (Invitrogen). The cancer cells were gated out with RFP (for luciferase reporter expression). The percentage of dead cells (SYTOX Green+) was calculated in FlowJo.

### Tumor-infiltrating lymphocyte phenotyping

Male CD45.1 C57BL/6 recipients bearing established subcutaneous male OVA-expressing CIN+ iT-KAP tumors received an adoptive transfer of male CD45.2 OT-1 CD8+ T cells (KDM5D-KD or WT control) on day 13 post-tumor inoculation. Tumors were harvested 16 days after T cell transfer for OT-1 state profiling and dissociated using the Mouse Tumor Dissociation kit (Miltenyi). Single-cell suspensions were stimulated with H2Kb/SIINFEKL tetramer (gifted from NIH Tetramer Core Facility at Emory, catalog no. 79539, 20 μg/mL) in T cell culture media supplemented with 1 × protein transport inhibitor cocktail (eBioscience, catalog no. 00-4980-93) for 3 h at 37 °C, then washed twice in PBS. Cells were then stained with TruStain FcX Plus anti-mouse CD16/32 antibody (BioLegend, catalog no. 156604, 1:50) and Ghost Dye UV450 viability stain (TONBO, catalog no. 13-0868-T100, 1:1000), incubated for 30 min at 4 °C in the dark, and washed once in PBS. Surface staining was performed according to the BioLegend surface staining protocol using CD8-BUV615 (BD, catalog no. 613004), CD45.2-BUV805 (BD, catalog no. 569200), LAG3-BV785 (BioLegend, catalog no. 125219), PD1-APC/Fire810 (BioLegend, catalog no. 135252), CD3e-PE/Cy5.5 (Thermo, catalog no. 35-0031-80), TIM3-APC/Fire750 (BioLegend, catalog no. 134017), followed by fixation, permeabilization, and intracellular staining using the eBioscience FoxP3/Transcription Factor Staining kit (Invitrogen, catalog no. 00-5523-00) per the manufacturer’s instructions with antibodies including Perforin-Pacific Blue (BioLegend, catalog no. 154408), TOX-AF647 (BD, catalog no. 568356), and Granzyme B-PE/Dazzle594 (BioLegend, catalog no. 372216). Single-color compensation samples were prepared with UltraComp eBeads Plus (Invitrogen, catalog no. 01-3333-42). Samples were acquired on a Cytek Aurora spectral flow cytometer and analyzed in FlowJo.

### RNA isolation and real-time quantitative PCR

RNA from cell lines was isolated using the RNeasy Kit (Qiagen) and TRIzol. RNA was reverse transcribed using a 5× All-In-One MasterMix (Applied Biological Materials Inc.). Real-time qPCR was performed using SYBR Green PCR Master Mix (Applied Biosystems) using the 7500 Fast Real-Time PCR System (Applied Biosystems). PCR product specificity was confirmed with a melting-curve analysis. Relative messenger RNA expression was calculated using the 2-ΔΔCt method. Relative expression was calculated by normalizing to Rn18s. The primer sequences are: Mouse *Kdm5d* forward (F): GCTGCTTGTGGTTGGAATCTC; reverse (R): AATGCTGAAAACACCATGCCC; *Rn18s* F: GTAACCCGTTGAACCCCATT; R: CCATCCAATCGGTAGTAGCG; *Tox* F: TGCCATTAAGGGCCAGAATCC; R: TCAGGTATTCCTTCTTTGCAGC; *Lag3* F: CTGGGACTGCTTTGGGAAG; R: GGTTGATGTTGCCAGATAACCC; *Trbc1/2* (B6) F: CAGGCCTACAAGGAGAGCAA; R: CTGAAAGCCCATGGAACTGC; *Tcrb* transgene (OT-1) F: ACGTGTATTCCCATCTCTGG; R: CTGTTCATAATTGGCCCCA; *Pdcd1* F: ACCCTGGTCATTCACTTGGG; R: CATTTGCTCCCTCTGACACTG; Spliced *Xbp1* F: AAGAACACGCTTGGGAATGC; R: CTGCACCTGCTGCGGAC; *Srebf2* F: GTTGACCACGCTGAAGACAGA; R: CACCAGGGTTGGCACTTGAA; *Havcr2* F: TCAGGTCTTACCCTCAACTGTG; R: GGGCAGATAGGCATTTTTACCA.

### RNAseq

RNA from cell lines was isolated using the RNeasy Kit (Qiagen). Each condition contained triplicate samples. RNAseq was performed by the Advanced Technology Genomics Core at The University of Texas MD Anderson Cancer Center. Libraries were generated using Illumina Stranded Total RNA Prep Ligation with Ribo-Zero Plus Kit following the manufacturer’s protocol. The NextSeq RTA (version 2.4.11.0) was used for base calling. Bcl2fastq (version 2.20.0) was used to convert the .bcl files into .fastq files. Fastq files were aligned to the mouse reference genome (mm10) using HISAT2 software and assembled using StringTie. R package DESeq2 was used for data normalization and differential expression analysis. The differentially expressed genes were analyzed using the Reactome database (https://reactome.org/) to identify the enriched pathways.

### CUT&RUN and analysis

CUT&RUN was performed by the Epigenomics Profiling Core at The University of Texas MD Anderson Cancer Center as described previously (Meers et al. 2019a) with some modifications. Briefly, nuclei isolated from ∼500,000 mouse OT-1 CD8+ T cells were immobilized on Concanavalin A-coated magnetic beads (Bangs Laboratories) and permeabilized using wash buffer containing 0.02% digitonin (Sigma). Bead-bound nuclei were incubated with rabbit IgG (Millipore), H3K4me1, H3K4me2, H3K27ac (all from Abcam), and H3K4me3 (EpiCypher) antibodies overnight at 4 °C. The next day, targeted chromatin digestion was achieved by pAG-MNase (EpiCypher) binding and incubation with CaCl_2_ at 4 °C, followed by CUT&RUN DNA purification using MinElute columns (Qiagen). Libraries from CUT&RUN DNA were prepared using NEBNext Ultra II DNA Library Prep Kit for Illumina (New England Biolabs) according to the manufacturer’s instructions. Sequencing was performed on an Illumina NextSeq 500 platform to obtain 5-10 million 75nt PE reads per sample. For analysis, reads were aligned to the mouse (mm9) and spike-in *E. coli* genome using Bowtie2 and unique read counts were recorded. The mouse and *E. coli* integrated genomes were built using Bowtie2. The scale factor was calculated as 1% of the integrated genome unique read counts divided by *E. coli* genome unique read counts. BigWig files were generated using bamCoverage with “—scaleFactor” set according to the above calculation and were visualized using IGV. SEACR (Meers et al. 2019b) was used for peak calling.

### Western blotting

For western blot analysis, cells were lysed on ice in RIPA buffer (Boston BioProducts) supplemented with protease and phosphatase inhibitors (Roche), then sonicated for 3 cycles (30 s on/30 s off) using a Biorupter (Diagenode). Protein concentrations were determined using the Pierce BCA Protein Assay kit (Thermo Fisher Scientific) on a CLARIOstar plate reader (BMG LABTECH). Samples were mixed with NuPAGE LDS Sample Buffer (4 ×; Invitrogen) containing 10% β-mercaptoethanol, boiled at 95 °C for 10 min, and loaded onto NuPAGE 4–12% Bis-Tris gels (Thermo Fisher Scientific) alongside Precision Plus Protein Dual Color Standards (Bio-Rad). The primary antibodies used include: anti-phos-LCK Y394 (BioLegend, catalog no. 933102, 1:1,000), anti-LCK (Cell Signaling Technology, catalog no. 2752, 1:1,000), anti-phos-ZAP70 Y319 (Cell Signaling Technology, catalog no. 2717, 1:1,000), anti-phos-ZAP70 Y493 (Cell Signaling Technology, catalog no. 2704, 1:1,000), anti-ZAP70 (Cell Signaling Technology, catalog no. 3165, 1:1,000), anti-phos-LAT Y132 (Invitrogen, catalog no. 44-224, 1:1,000), anti-LAT (Cell Signaling Technology, catalog no. 45533, 1:1,000), anti-phos-PLCγ1 Y783 (Cell Signaling Technology, catalog no. 14008, 1:1,000), anti-PLCγ1 (Cell Signaling Technology, catalog no. 5690, 1:1,000), anti-phos-ERK (Cell Signaling Technology, catalog no. 4370, 1:1,000), anti-ERK (Cell Signaling Technology, catalog no. 9102, 1:1,000), anti-Vinculin (Millipore, catalog no. 05-386, 1:500), anti-β-actin (Sigma, catalog no. A2228, 1:2,000), anti-TCR beta (Cell Signaling Technology, catalog no. 65123, 1:1,000). The secondary antibodies used include anti-rabbit IgG, HRP-linked (Cell Signaling Technology, catalog no. 7074, 1:5,000) and anti-mouse IgG, HRP-linked (Cell Signaling Technology, catalog no. 7076, 1:5,000).

To prepare restimulated CD8+ T cells for western blot, activated OT-1 CD8+ T cells were washed twice in plain HBSS and once in RPMI-1640 with 0.5% FBS, then starved for 2 h at 37 °C (2 × 10^6^/mL in RPMI-1640 with 0.5% FBS) to lower basal kinase activity. Cells were resuspended in pre-warmed HBSS containing Ca^2+^ and Mg^2+^, aliquoted at 1 × 10^6^ per condition into tubes held at 37 °C, stimulated with H2Kb/SIINFEKL tetramer (gifted from NIH Tetramer Core Facility at Emory, catalog no. 79539, 1 μg/mL) for 0, 30 s, 1, 2, and 5 min. Reactions were quenched with an equal volume of ice-cold 2 × RIPA buffer containing 2 × protease and phosphatase inhibitors.

### Amplex Red Cholesterol Assay

CD8+ T cells were collected with 1 × RIPA buffer and the protein concentration was measured with a plate reader as described above. Samples with the volume containing equal amount of protein were prepared and analyzed using the Amplex Red Cholesterol Assay Kit (Invitrogen) following the manufacturer’s protocol.

### In vitro chronic versus acute stimulation

Naïve CD8+ T cells were isolated from male mice and activated on day 0 with anti-CD3/CD28 Dynabeads in 30 U/ml IL-2-containing media. On day 3, beads were removed and cells were electroporated to knock down KDM5D, then allowed to recover overnight. On day 4, cells were divided into two arms: under acute conditions, cells were maintained in IL-2 without further TCR engagement through day 12; under chronic conditions, cells received repeated anti-CD3/CD28 restimulation on days 4, 6, and 9 to drive an exhaustion-like state. Both arms were harvested on day 12 for downstream analysis.

### In vitro statin treatment and cancer cell proliferation assay

CD8+ T cells were treated with lovastatin (Selleck, catalog no. S2061, prepared in DMSO) at the indicated dosage for 48 h and collected for qPCR analysis. Cancer cells (1 × 10^4^) were plated in 96-well plates and treated with lovastatin at the indicated dosage for 72 h. Cells were incubated in IncuCyte (Sartorius) and images were taken every 8 h for 72 h. Cell confluency was calculated in the IncuCyte software ZOOM 2018A.

### In vivo lovastatin and CD8 neutralizing antibody treatment

Lovastatin was prepared in 6% PEG400, 1% Propylene glycol, 0.1% Tween-80, as described previously (Fernandez et al. 2020). Male C57BL/6 mice bearing CIN+ iT-KAP subcutaneous tumors were treated with 60 mg/kg lovastatin or vehicle by oral gavage once daily. Tumor sizes and body weights were measured twice a week. One shot of 500 ug per mouse anti-mouse CD8β (BioXCell, catalog no. BE0223, Clone 53-5.8) was given the day before treatment started and then every two weeks.

### Immunofluorescence staining of mouse tumors

Mouse tumors were fixed in 10% neutral-buffered formalin, paraffin-embedded, and sectioned at 5 μm thickness. Sections were deparaffinized in xylene, rehydrated through a graded ethanol series, and rinsed in distilled water. Antigen retrieval was performed by heating the sections in citrate-based antigen retrieval buffer (pH 6.0) using a pressure cooker. Endogenous peroxidase activity was quenched by incubating the sections with 3% hydrogen peroxide for 10 min at room temperature (RT), followed by a wash with Tris-buffered saline containing 0.05% Tween-20 (TBST). Sections were blocked with 10% normal horse serum for 1 h at RT and incubated overnight at 4 °C with anti-CD8α primary antibody (Cell Signaling Technology, catalog no. 98941). After three washes with TBST, sections were incubated with an HRP-conjugated secondary antibody for 1 h at RT. Following an additional three TBST washes, signal was developed using Alexa Fluor 488-labeled tyramide conjugates (Invitrogen, catalog no. B40943). Sections were counterstained with DAPI and mounted using VECTASHIELD Antifade Mounting Medium (Vector Laboratories, catalog no. H-1000-10).

### ScRNAseq data analysis

CD8+ T cells were extracted from the all-cell annotated data and were processed using R package Seurat v5.0.1 (Hao et al. 2024). Specifically, the expression data went through SCTransform normalization, PCA, batch effects correction using Harmony v.1.2.0 (Korsunsky et al. 2019), UMAP, clustering, and differential expression (DE) analysis to detect top genes in different clusters. Marker genes were visualized by dot plots using R package ggplot2 v3.5.2 and top genes in clusters from DE helped with annotating CD8+ T cell subtypes. The numbers of cells in CD8+ T cell subtypes across sexes were extracted, based on which their percentages were calculated. Statistical significance was tested for the percentages using R package stats. GSEA analysis was conducted using R package clusterProfiler v4.10.0 (Wu et al. 2021) for the HALLMARK pathways based on fold-change pre-ranked genes from the DE results. GSVA scores of the CD8+ T cells were calculated for the HALLMARK_CHOLESTEROL_HOMEOSTASIS pathway using R package GSVA v1.50.0 (Hanzelmann et al. 2013). The boxplot showing GSVA scores in the three CD8+ T cell subtypes was generated using R package ggpubr v0.6.0.

All human studies were performed in accordance with the Declaration of Helsinki and approved by the Institutional Review Board of The University of Texas MD Anderson Cancer Center (protocol number: LAB09-0373; principal investigator: Dr. Scott Kopetz). Human tumor specimens were collected after obtaining written informed consent from all participants.

### Statistical analysis

All statistical tests were performed in GraphPad Prism v.10 and R, and the statistical analyses are noted in figure legends.

## Data availability

The mouse scRNAseq data have been deposited in GEO previously (GSE229559) (Hsu et al. 2023). The human scRNAseq data and clinical information are available upon request. The bulk RNAseq and CUT&RUN data from mouse CD8+ T cells were deposited in GEO (GSE330655 and GSE331334).

## Competing interest statement

R.A.D. is a founder and advisor of Tvardi Therapeutics, Inc. and Regenimo Biosciences, Inc., an advisor for Modi Ventures, LLC, and a founder and equity holder for Sporos Bioventures, LLC. The work of this study bears no direct relevance to these entities. J.P.S. has received grants, research support and/or has research collaborations with Natera, Inc., BostonGene, Revolution Medicine, and Summit Therapeutics. J.P.S. has served as a consultant for and/or holds equity in Engine Biosciences and NaDeNo Nanoscience. The rest authors declare no competing interests.

## Acknowledgements

We thank Dr. James Allison for insightful suggestions and criticisms and all members of the Allison laboratory for helpful discussions and support. We thank Dr. Yejing Ge for the input on the epigenetic study. J.Li was supported by the Odyssey Fellowship Program of the University of Texas MD Anderson Cancer Center and the Crum Foundation. A.B.S. was supported by the Triumph postdoctoral training program at MD Anderson supported by the Cancer Prevention & Research Institute of Texas (CPRIT) (grant no. RP210028). J.Liu and S.E.W. were supported by The University of Texas MD Anderson Cancer Center Moon Shots Program. W.H.W. and M.S. were supported by the CPRIT Training Program (RP210028). A.K.J. was partially supported by institutional support to the Epigenomics Profiling Core (EpiCore) at The University of Texas MD Anderson Cancer Center. J.P.S. was supported by the Col. Daniel Connelly Memorial Fund, the Andrew Sabin Family Fellowship Award as an Andrew Sabin Family Foundation Fellow at The University of Texas MD Anderson Cancer Center, CPRIT (RR180035 & RP240392) as a CPRIT Scholar in Cancer Research, and Conquer Cancer (2022CDA-7604125121). The human scRNAseq study was supported by Celsius Therapeutics. S.K. was supported by the Cancer Center Support Grant National Cancer Institute (NCI) P30CA016672. Work in R.A.D.’s laboratory was supported by NIH R01 CA231360, CPRIT (RP220364), and the Harry Graves Burkhart III Distinguished University Chair in Cancer Research.

The MDACC Advanced Cytometry & Sorting Core Facility was supported by NCI P30CA016672. Sequencing data were generated in part through the use of the Advanced Technology Genomics Core (ATGC) at The University of Texas MD Anderson Cancer Center, which receives partial support from the NCI under grant P30CA016672. The research reported in this publication was not directly funded through the P30CA016672 grant to The University of Texas MD Anderson Cancer Center and is not within the scope of such grant.

## Author Contributions

J.Li and R.A.D. designed the project, conceived the experiments, interpreted the observations, analyzed the data, and wrote the manuscript. C.Y.C. assisted with in vitro assays and mouse experiments. A.B.S. performed the tumor penetrance assay. A.B.S. and P.T.D.L. provided reagents and conceptual and technical support for tumor dissociation, flow cytometry and in vivo mouse experiments. J.Liu performed bioinformatic analyses. S.J. performed the in vivo treatment and maintained, monitored and recorded the health condition of mice. C.L. provided CIN+ iT-KAP organoids. Z.Z., W.H.W, M.S., X.W. and R.C.M. assisted with the in vitro assays and mouse experiments. A.K.J. provided CUT&RUN service and guidance on data analysis. N.H., F.Z. and M.Z. performed analyses on patient scRNAseq data. S.E.W. provided guidance and assisted with scRNAseq analyses. N.R.F. provided guidance and assisted with cholesterol metabolic analyses. D.J.S. edited the manuscript and reviewed data. J.P.S. and S.K. provided patient scRNAseq data. All authors edited and approved the final version of the manuscript.

## Notes

### Summary of Updates

Supporting Information (Figures S1 to S6) have been added.

