## Supporting Information Figs S1 to S6. for "The Y chromosome gene KDM5D restrains CD8+ T cell antitumor immunity through TCR and cholesterol-exhaustion programs"

PNAS

**This PDF file includes:**

Figures S1 to S6

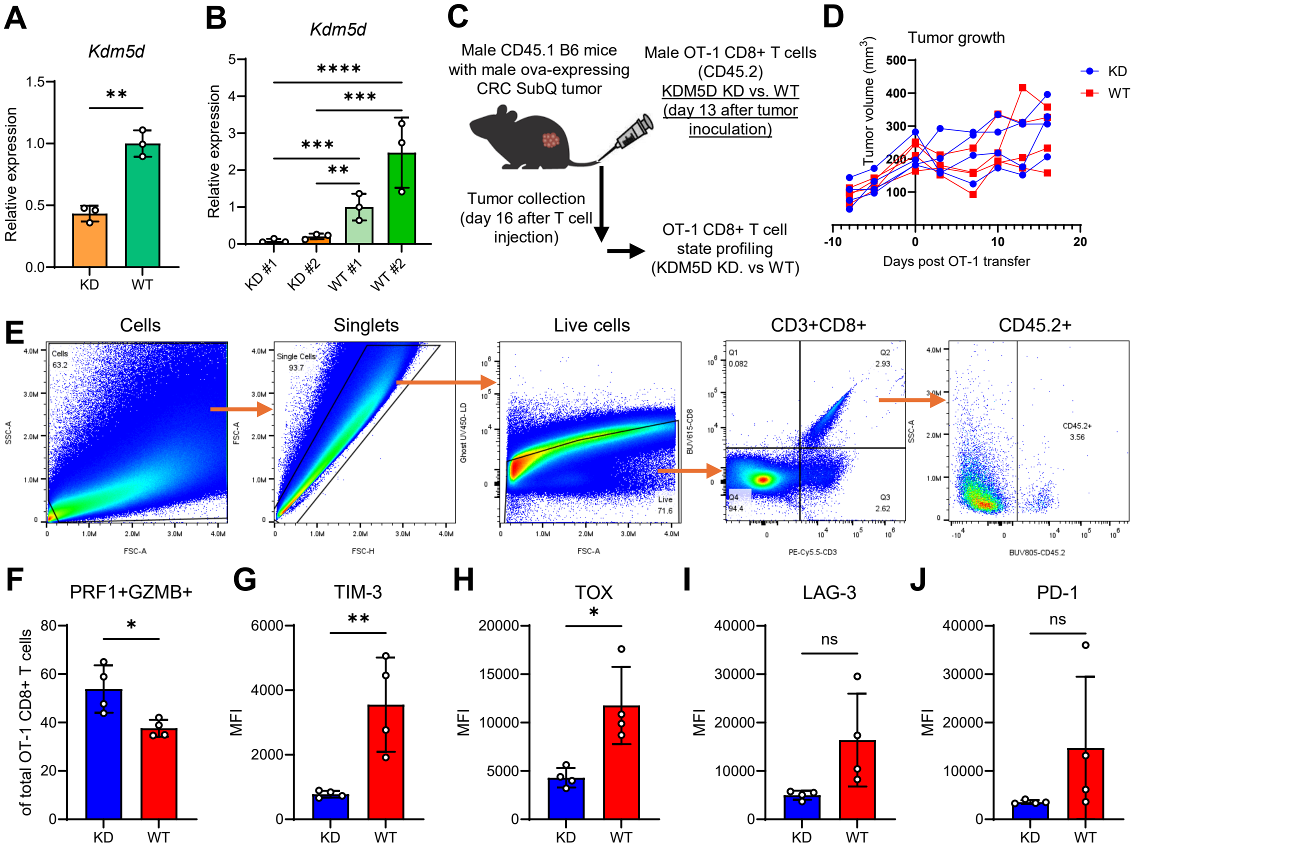


**Supplemental Fig. S1. KDM5D contributes to the suppression of CD8+ T cell anti-tumor function.**

**A**. qPCR of KDM5D in male OT-1 CD8+ T cells. **B**. qPCR of KDM5D in tumor-infiltrating CD8+ T cells of CMT93-KRAS-G12D-OVA tumors. **C**. Experimental design. **D**. SubQ tumor growth curve. **E**. Flow cytometry gating strategy. **F**. Percentage of indicated OT-1 CD8+ T cell subpopulation in all OT-1 CD8+ T cells. **G-J**. Expression levels of the indicated marker in all CD8+ T cells. All bar graphs show mean value ± s.d.; *P* was derived with one-way ANOVA with multiple comparisons (**B**) and two-tailed unpaired t-test (**A**, **F-J**).

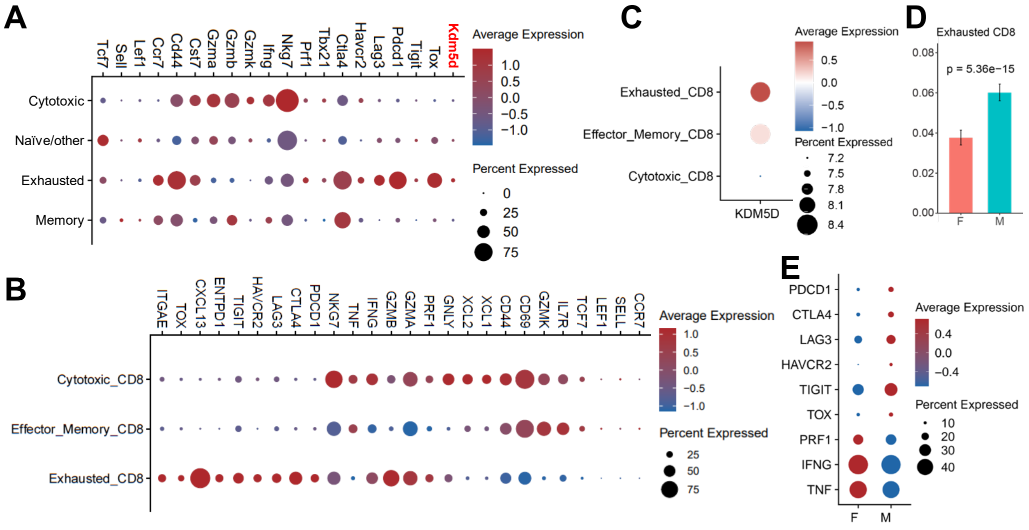


**Supplemental Fig. S2. Analyses of scRNAseq of mouse and human CRC.** **A**. Expression of KDM5D and markers used to define CD8+ T cell subtypes in the mouse scRNAseq dataset (male only). **B**. Genes used to annotate CD8+ T cell subtypes in human scRNAseq dataset (male and female combined). **C**. KDM5D expression in patient scRNAseq (male only). **D**. Percentage of exhausted CD8+ T cells in all CD8+ T cells (Pearson's chi-squared test). **E**. Expression level of the indicated genes in patient scRNAseq.

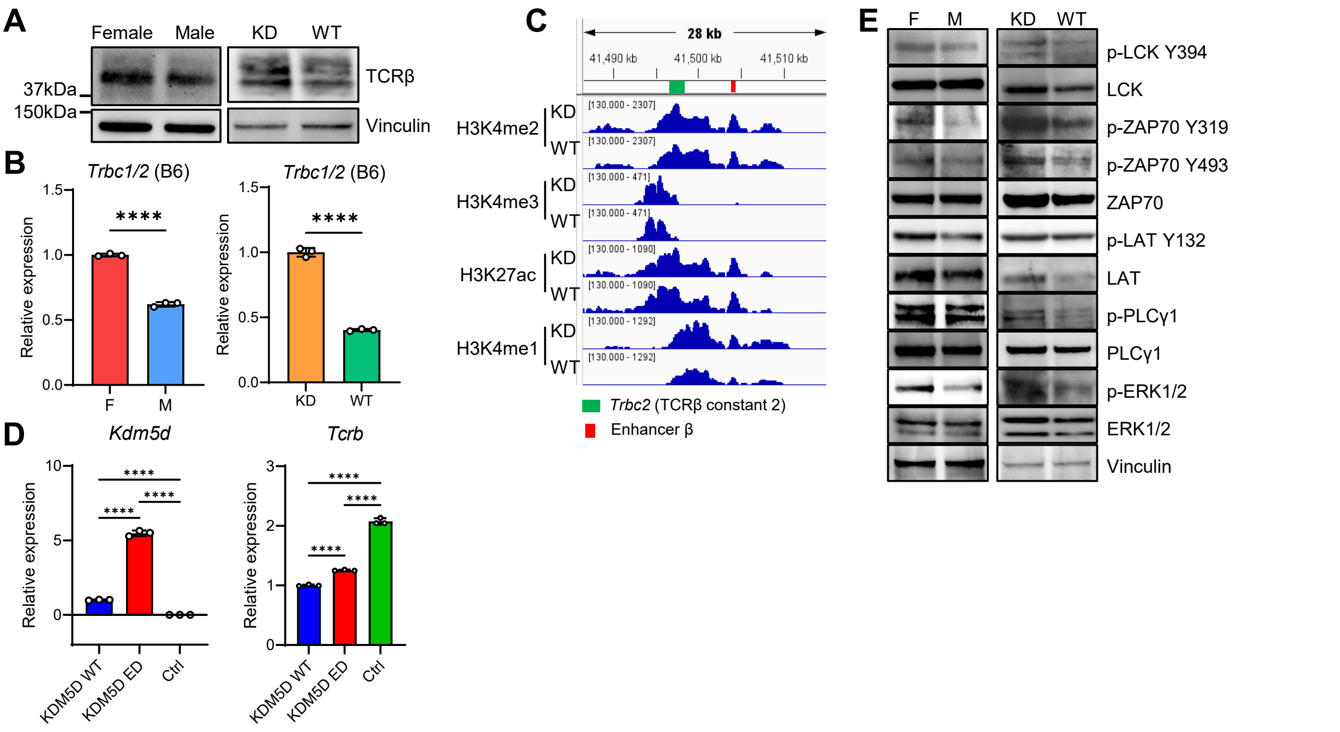


**Supplemental Fig. S3. KDM5D restrains TCR abundance and proximal signaling via an epigenetic mechanism.**
**A**. WB of total TCRβ and vinculin in OT-1 T cells. **B**. qPCR of T Cell Receptor Beta Constant (*Trbc*) 1/2 gene in B6 CD8+ T cells. **C**. CUT&RUN track at TCRβ gene locus. **D**. qPCR of female CD8+ T cells overexpressing mouse WT, ED (enzymatic-dead) KDM5D, or control vector. **E**. WB of TCR signaling markers. For **B** and **D**, data are mean value ± s.d.; *P* was derived with two-tailed unpaired t-test (**B**) and one-way ANOVA with multiple comparisons (**D**).

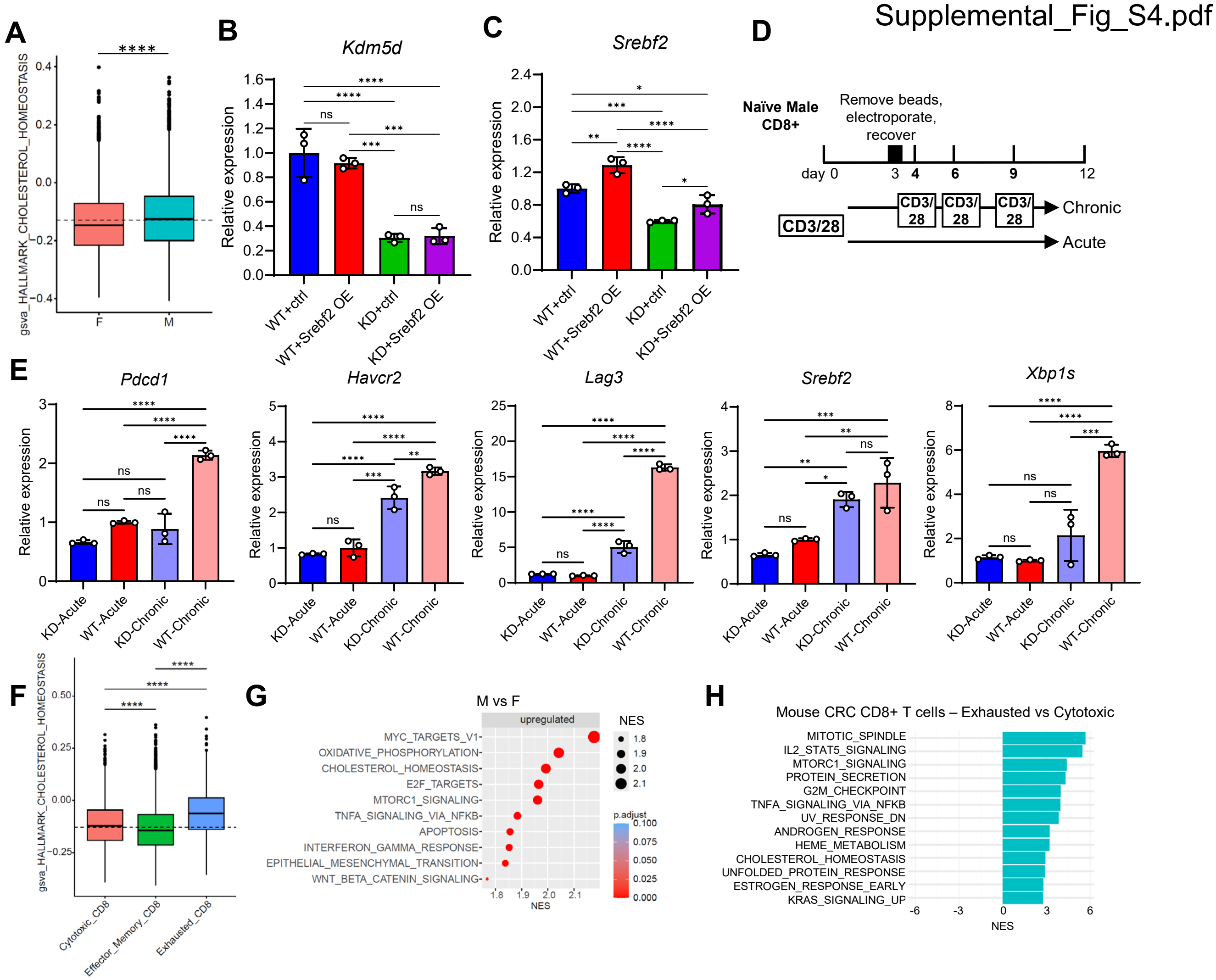


**Supplemental Fig. S4. KDM5D contributes to sex differences in cholesterol homeostasis and exhaustion-associated programs in male CD8+ T cells.**

**A**. GSVA analysis (Wilcoxon rank sum test). **B**, **C**. qPCR of *Kdm5d* and *Srebf2* in male T cells with WT or KD of KDM5D and OE of SREBF2 or vector control. **D**. Approach for generating exhausted CD8+ T cells in vitro. **E**. qPCR of genes in indicated conditions. **F**. GSVA analysis (Wilcoxon rank sum test). **G**. Differential Hallmark pathways in male versus female exhausted CD8+ T cells in patient scRNAseq. **H**. Differential Hallmark pathways in exhausted versus cytotoxic CD8+ T cells in iKAP tumor scRNAseq (all males). For **B**, **C**, and **E**, data are mean value ± s.d.; *P* was derived with one-way ANOVA with multiple comparisons. NES: normalized enrichment score.

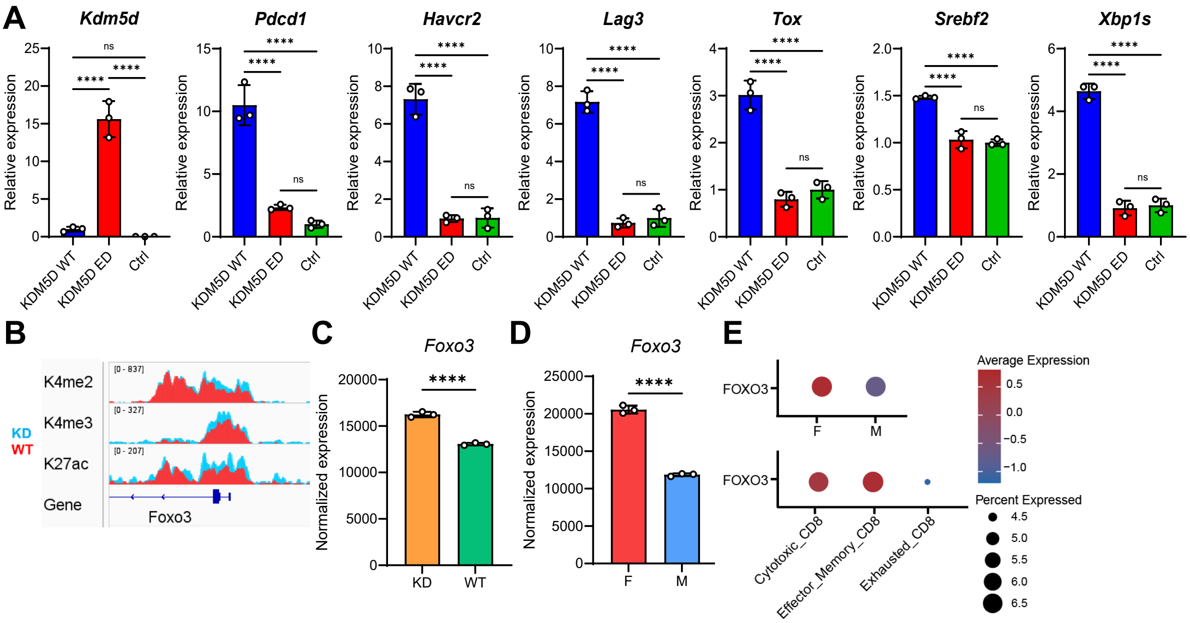


**Supplemental Fig. S5. KDM5D may epigenetically regulate *Foxo3* expression in male CD8+ T cells.**

**A**. qPCR of female CD8+ T cells overexpressing mouse WT, ED KDM5D or control vector. **B**. CUT&RUN track at *Foxo3* gene locus. **C**, **D**. Normalized expression of *Foxo3* in indicated cells from RNAseq data. **E**. *FOXO3* expression in patient scRNAseq. For **A**, **C** and **D**, data are mean value ± s.d.; *P* was derived with one-way ANOVA with multiple comparisons (**A**) and two-tailed unpaired t-test (**C** and **D**).

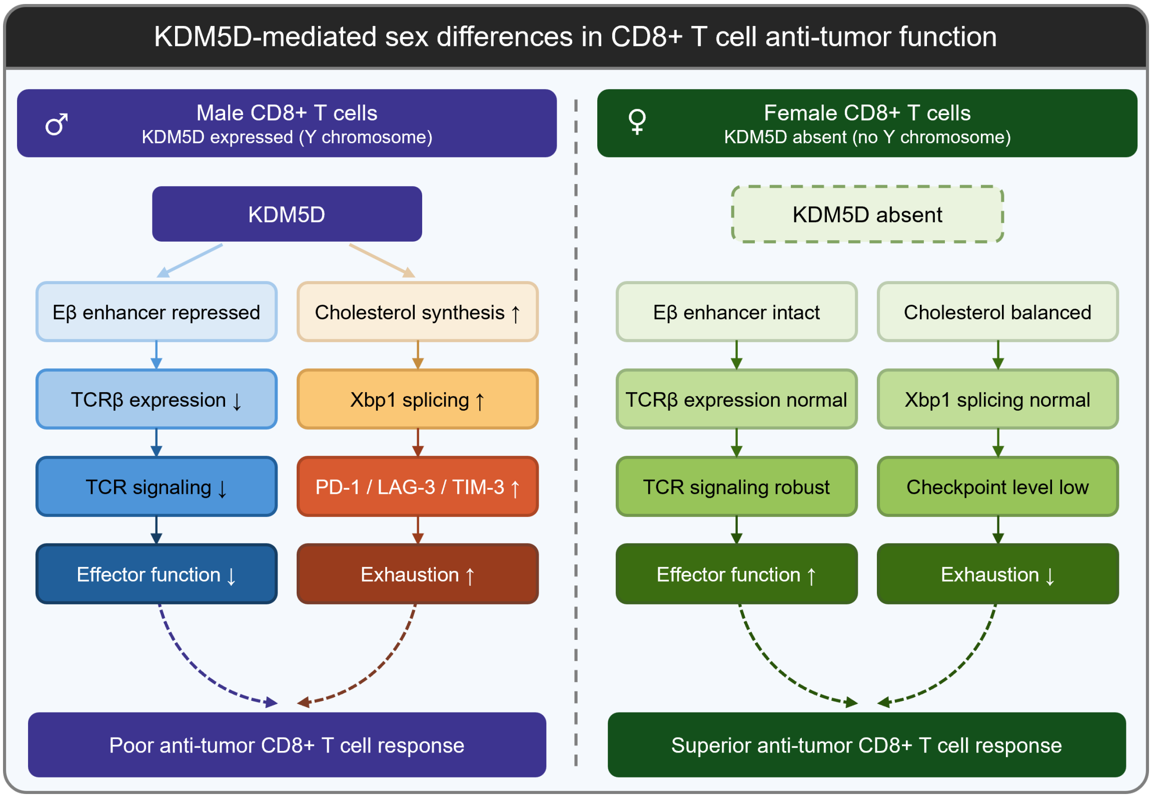


**Supplemental Fig. S6. Schematic representation of KDM5D-induced sex differences in CD8+ T cell function.**
